# Single cell transcriptomics reveal mast cell heterogeneity in the human gut

**DOI:** 10.1101/2024.11.29.626054

**Authors:** Hind Hussein, Elodie Modave, Nathalie Stakenborg, Marcello Delfini, Katy Vandereyken, Alejandro Sifrim, Thierry Voet, Guy E. Boeckxstaens

**Affiliations:** Center for Intestinal Neuroimmune Interactions, Translational Research Center for Gastrointestinal Disorders (TARGID), KU Leuven Department of Chronic Diseases, Metabolism and Ageing (CHROMETA), KU Leuven, Leuven, 3000, Belgium; Laboratory of Tumor Inflammation and Angiogenesis, Center for Cancer Biology, VIB, Leuven, Belgium; Laboratory of Tumor Inflammation and Angiogenesis, Center for Cancer Biology, Department of Oncology, KU Leuven, Leuven, Belgium; Laboratory of Reproductive Genomics, Department of Human Genetics, KU Leuven, Leuven, Belgium; KU Leuven Institute for Single Cell Omics, KU Leuven, Leuven, Belgium; Laboratory of Multi-Omic Integrative Bioinformatics, Department of Genetics, KU Leuven, Leuven, Belgium; Leuven AI Institute, KU Leuven, Leuven, Belgium

## Abstract

Gut mast cells are key players in infection and inflammation, as well as in homeostasis. Mast cells regulate gastrointestinal (GI) function by controlling vascular and epithelial permeability, and interacting with the immune system and the enteric nervous system. Mucosal (MC_T_) and connective tissue (MC_TC_) mast cells coexist in the gastrointestinal tract, but are located in distinct microanatomical niches. However, little is known about the transcriptional heterogeneity of human intestinal mast cells. Additionally, whether distinct microanatomical niches instruct mast cell phenotypes in the human gut is currently unknown. We therefore performed 10x single cell RNA sequencing on healthy rectal biopsies and on dissected layers of human sigmoid colon. We identified five transcriptionally distinct mast cell subsets (MC1-5) which exhibited a layer-specific distribution in the healthy human colon and distinct cytokine, chemokine, protease, and transcription factor profiles, and were associated with putative immune and neuro-immune functions. This study provides a framework for the integration of mast cell heterogeneity in the study of gastrointestinal disease mechanisms in humans.

## Introduction

Mast cells are tissue-resident cells found in barrier tissues and exerting a wide range of functions^3^. While most commonly associated with allergy and the onset of anaphylaxis, mast cells are ancient cells that have evolved long before the adaptive immune system^4,5^. Mast cells are morphologically distinct among immune cells and can be identified by their electron-dense secretory granules which contain preformed mediators, most notably serine proteases, histamine, TNF-α, and serotonin^6^. Upon activation, mast cells release their granule contents into the extracellular environment and initiate *de novo* synthesis of cytokines, chemokines, and eicosanoids. Mast cell activation is classically triggered by IgE antibodies, but can also be mediated by complement activation, pattern recognition receptors, or occur following tissue damage and in response to neuronal peptides. Recent work has emphasized the role of Mas-related G protein–coupled receptor X2 (MRGPRX2) in IgE-independent, drug-induced pseudoallergic mast cell reactions^3^. Aside from drugs, endogenous antimicrobial peptides and neuropeptides also trigger MRGPRX2-mediated mast cell activation. This testifies to the privileged role mast cells hold as intermediaries between the nervous and immune systems, which places mast cells as a coordinator of neuroimmune reactions at the tissue level. Overall, the breadth of mast cell activation signals is key to the sentinel function exerted by mast cells in barrier tissues.

Mast cells have been histologically classified based on granule protease expression into two subsets. Mucosal mast cells, or MC_T_ in humans, express tryptase, while connective tissue mast cells, or MC_TC_ in humans, co-express tryptase and chymase. This distinction also exists in mice, where protease expression patterns differ, with mucosal mast cells expressing mast cell proteases 1 (Mcpt1) and 2 (Mcpt2), and connective tissue mast cells expressing Mcpt4-7. Interestingly, expression of Mrgprb2, the murine ortholog of MRGPRX2, has been tied to the connective tissue mast cell phenotype, while mucosal mast cells are negative for Mrgprb2 expression^7^. This binary classification of mast cells also accounts for the heterogeneity of developmental origins of mast cells. Murine fate-mapping experiments have shown that mucosal mast cells, arising from bone-marrow progenitors, migrate to the intestine and the lung in a T cell-dependent manner, while connective tissue mast cells are long-lived cells which develop from fetal liver progenitors during embryogenesis^8^.

Gut mast cells play a variety of roles in infection^9^ and inflammation^1,2^, but are also essential to regulate intestinal homeostasis. Mast cells notably control vascular and epithelial permeability in the intestine via the production of histamine and proteases^10,11^. Mast cells also support intestinal IgA responses, essential for intestinal tolerance and controlling microbiota composition, by participating in the organization of the Peyer’s patch niche, through the recruitment of GPR35^+^ cDC2s^12^. Mast cells also collaborate with the enteric nervous system to regulate sensation, secretion, and motility in the digestive tract^13^. Interestingly, both histologically-defined mast cell subsets coexist in the gastrointestinal tract, but are located in distinct microanatomical niches^11^. The gut wall is composed of three layers which are dedicated to different facets of gastrointestinal function. The mucosa is composed of the intestinal epithelium and the lamina propria and is involved in immune defense. The submucosa, involved in secretion and sensation, contains blood and lymphatic vessels, glands, and a nervous plexus, the submucosal plexus. The muscularis controls intestinal peristalsis and is formed of two muscle layers surrounding the myenteric plexus^14^. Mucosal mast cells reside in the mucosa, while connective tissue mast cells were shown to be enriched in the muscularis. Incidentally, Mrgprb2-expressing mast cells were also shown to be enriched in the intestinal muscularis while mast cells in the mucosa were mostly negative for Mrgprb2^7^.

The advent of single-cell sequencing has challenged the dichotomic characterization of mast cells based on protease expression and localization within connective or mucosal tissues and recently revealed a greater heterogeneity of mast cells across human organs^7,15^. To date, the heterogeneity of human mast cells within intestinal microanatomical niches has not been appraised. Here, we profiled mast cells in the healthy human sigmoid colon and rectum using single cell transcriptomics. We identified five transcriptionally distinct mast cell subsets which exhibited a layer-specific distribution in the healthy human colon and distinct cytokine, chemokine, protease, and transcription factor profiles, and were associated with putative immune and neuro-immune functions.

## Materials and methods

### Human samples

#### Ethics

All participants gave written informed consent and the Medical Ethical Committee of UZ/KU Leuven approved the protocols (ethical approval number S62059).

#### Colonic resection tissue

Colon tissues were obtained from 3 patients (1 F and 2 M, median age of 58 years IQR [55.5-63]) undergoing left hemicolectomy for colonic carcinoma.

#### Rectal biopsies

Healthy volunteers (HV) free of gastrointestinal symptoms, with no history of gastrointestinal disease or surgery were recruited by public advertisement. IBS patients fulfilling the Rome IV criteria were recruited at the outpatient clinic of the University Hospitals Leuven. 5 HV (3 F and 2 M, median age of 35 years IQR [23-37]) were invited to undergo a proctoscopy to collect rectal biopsies.

### Reagents and media

Information on reagents and media can be found in **Table S1**.

### Single cell suspension from gut tissue

Colon tissue was dissected into layers. Mucosal layers (colon mucosa and rectal biopsies) were incubated at 37°C under rotation in epithelial removal buffer for 15 min with dithiothreitol and 15 min without to remove epithelial cells. Samples were minced and incubated 30 min at 37°C under rotation in digestion medium with 20U/mL DNase I and 0.4mg/mL liberase (TH for muscularis, TM for mucosa and submucosa) or 0.1mg/mL liberase TM for rectal biopsies. Samples were resuspended in FACS buffer after red blood cell lysis.

### Fluorescence-associated cell sorting

Single cell suspensions were stained in FACS buffer containing FcR blocking antibody and anti-CD45 FITC and incubated for 25 min at 4°C in the dark. Dead cells were excluded by staining with 7-aminoactinomycin D. Acquisition was performed on a Sony MA 9000 cell sorter with a 100µm nozzle. Live CD45^+^ cells were sorted (4.10^4^-10^5^ cells per patient).

### scRNA-seq library preparation and sequencing

Complementary DNA libraries were generated using the 10x Genomics Chromium Single Cell 3’ kit according to manufacturer’s instructions. Two lanes of 10,000 cells were loaded per sample. Sequencing was conducted with a NovaSeq6000 (Illumina) to a depth of at least 25,000 reads per cell.

### scRNA-seq analysis

Raw sequencing data were demultiplexed to generate fastq files, which were aligned to the human refence genome build GRCh38 (GENCODE v32/Ensembl 98), using STAR, with the CellRanger count function (10x Genomics – version 6.1.2). A single-cell expression matrix (i.e. counts by cells) was then generated by CellRanger and the generated data for each sample was uploaded to Seurat. SoupX^16^ was implemented on each sample (18 samples representing 8 patients: 5 rectal biopsies and 3 colonic resection tissues) to remove the ambient RNA by estimating the proportion of ambient RNA in each gene expression profile. SoupX compares the RNA distribution in low-expressing cells, where ambient RNA has a stronger influence, to high-expressing cells, where endogenous RNA dominates. Once the ambient RNA contribution is quantified, SoupX adjusts the data by subtracting this contamination, allowing for more accurate representation of true cellular RNA levels. Following a quality check and filtering of these samples (features: 400-6,000; mitochondrial gene content < 20%: UMIs: 1,000-50,000), cells were normalized, scaled and integrated using Harmony^17^. A total of 139,057 cells were kept. The highly variable genes of the integrated dataset (i.e. standard 2,000 genes) were then used for a dimensional reduction to plot each single cell on a 2-D Uniform Manifold Approximation and Projection (UMAP). Doublets were detected with the DoubletFinder algorithm^18^ where artificial doublet cells are generated for each sample and Euclidian distances between each real cell and artificial cells are calculated. If a real cell is close to artificial doublets, it is considered as a doublet. We investigated the doublet load (a doublet is created when multiple cells are captured in a single gel bead). For this, we set a number of generated artificial doublets (i.e. pN) of 25% of the merged real-artificial data (default value) and PC neighborhood size used to compute the proportion of nearest neighbors (i.e. pK) at 1% of the merged real-artificial data.

To remove dead cells, we visually selected the clusters that had a combination of high level of mitochondrial content and a low proportion of ribosomal content. We custom-set the thresholds using the median value of both metrics. After thoroughly cleaning the data, we were left with 137,087 cells. The differential gene expression between clusters was calculated using the FindAllMarkers function of Seurat with the Wilcoxon rank sum test to measure the level of expression of genes between different clusters. At least 10% of the cells in one cluster needed to express a gene at a minimum of 0.25 log_2_FC. Subsequently, volcano plots were generated.

Depending on their similar gene expressions, cells clustered in different groups (i.e. cell populations) after creation of a vector of resolutions.

pySCENIC (Single-Cell Regulatory Network Inference and Clustering)^19^ is a computational method that reconstructs Gene Regulatory Networks (GRNs) and determines cell states from single-cell RNA-seq data. By identifying key transcription factors that regulate these networks, pySCENIC provides insight into how these regulators influence the expression of downstream target genes. pySCENIC was utilized to identify transcription factors and evaluate their role in modulating gene expression in healthy samples. Its capacity to reveal transcription factor-target gene interactions in an unbiased manner at single-cell resolution makes it an essential tool for investigating the regulatory processes that drive cellular heterogeneity and dynamics.

gprofiler2 (v0.2.3)^20^ was used for pathway enrichment analysis (**Fig. 2D**) and is designed to interpret gene lists by mapping them to known biological pathways. It operates by taking the list of differentially expressed genes between clusters and performing an over-representation analysis (ORA), comparing the input list to a background dataset. It then identifies which biological pathways are statistically enriched in the gene list, meaning these terms are associated with more genes than would be expected by chance. gprofiler2 integrates various data sources, including KEGG, Reactome, and GO, to provide comprehensive coverage of biological processes, molecular functions, cellular components, and pathways.

**Figure 1.**
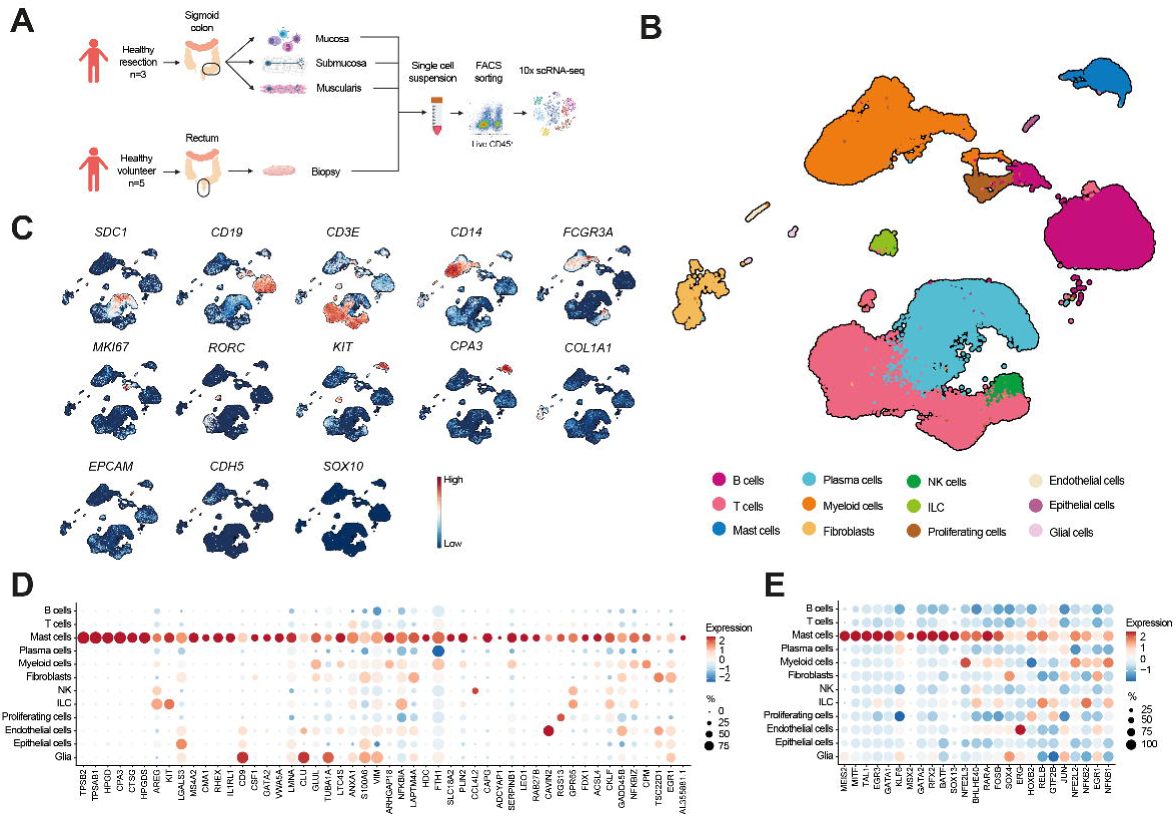
Single cell RNA sequencing of healthy human colonic and rectal immune cells. (A) Experimental design used in this study. (B) Annotated uniform manifold approximation and projection (UMAP) dimensionality reduction and clustering analysis of 10x scRNA-seq data obtained from sorted live CD45+ cells from rectal biopsies of 5 healthy volunteers and from dissected mucosa, submucosa, and muscularis externa layers of the healthy sigmoid colon of 3 hemicolectomy resection patients. (C) Feature plots showing the expression of *SDC1, CD19, CD3E, CD14, FCGR3A, MKI67, RORC, KIT, CPA3, COL1A1, EPCAM, CDH5,* and *SOX10* used for cell subset annotation at the CD45 level. (D) Dotplot showing the expression of the top 50 mast cell genes in the identified immune and non-immune cell subsets in healthy colon and rectum. (E) Dotplot showing the expression of the 25 top mast cell regulons identified via single-cell regulatory network inference and clustering (SCENIC) analysis in the identified immune and non-immune cell subsets in healthy colon and rectum. In dotplots, the color of dots represents expression level, while the size of dots represents the percentage of cells expressing the gene identified.

**Figure 2.**
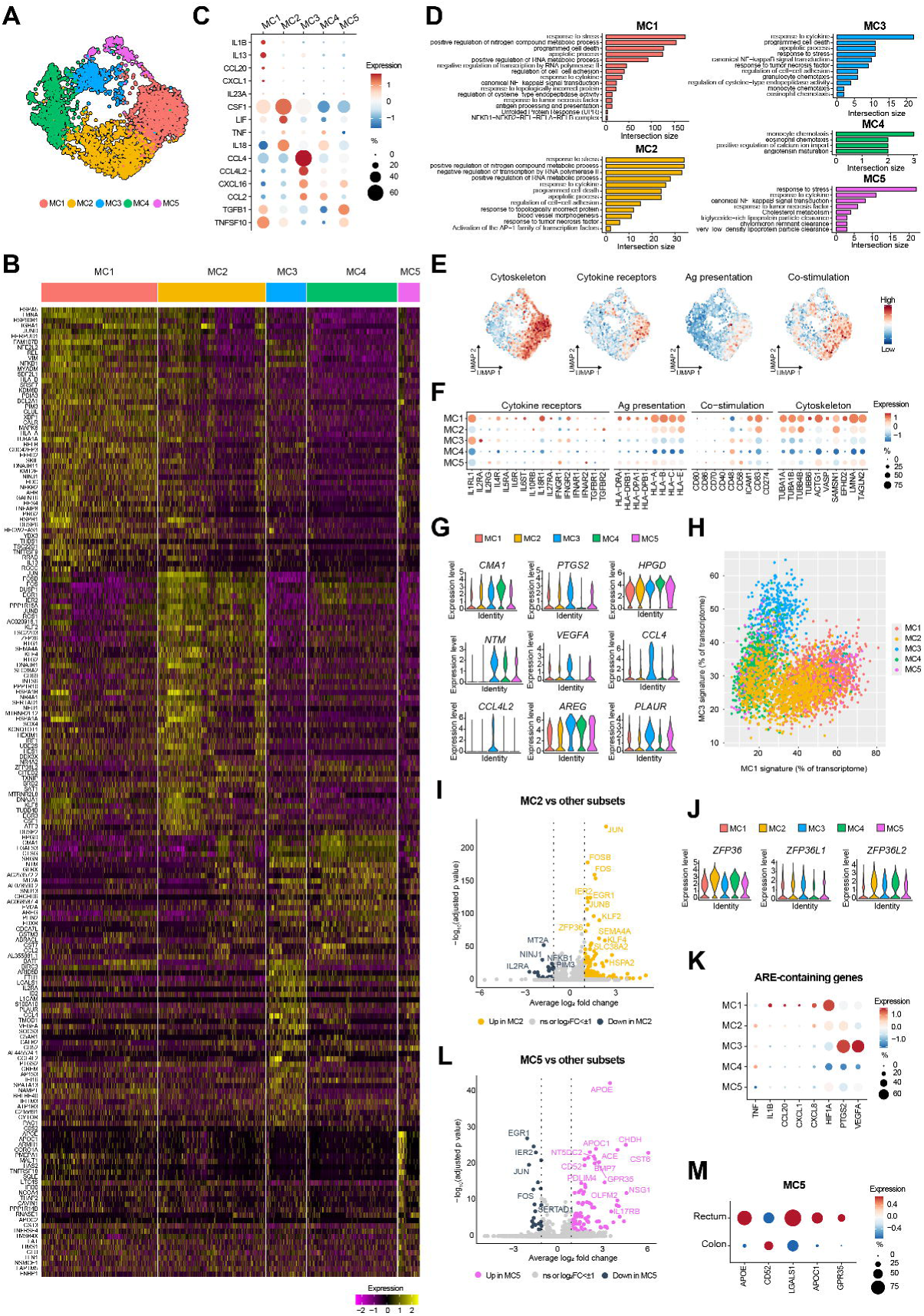
The human gut harbors five transcriptionally distinct mast cell subsets. (A) Annotated UMAP of mast cells from the healthy human colon and rectum. (B) Heatmap showing the top 50 genes expressed in human gut mast cell subsets. (C) Dotplot showing the expression of cytokine and chemokine genes in human gut mast cell subsets. (D) Pathway analysis in mast cell subsets. (E, F) Feature plots showing aggregated gene signature expression (E) and dotplot showing individual gene expression (F) of cytoskeleton, cytokine receptors, antigen presentation, and co-stimulation molecule genes in mast cell subsets. (G) Violin plots showing expression distribution of individual genes in mast cell subsets. (H) Per-cell expression of MC1 and MC3 signatures across all clusters. (I) Volcano plot showing differentially expressed genes in MC2 compared to other mast cells. (J) Violin plots showing expression distribution of tristetraprolin family genes in mast cell subsets. (K) Dotplot showing the expression of AU-rich elements (ARE)-containing genes in mast cell subsets. (L) Volcano plot showing differentially expressed genes in MC5 compared to other mast cells. (M) Dotplot comparing the expression of individual genes in MC5 from the rectum or the colon. In dotplots, the color of dots represents expression level, while the size of dots represents the percentage of cells expressing the gene identified.

The data used for the scatter plots in **Fig. 2H** was generated using UCell^21^, an R package designed to score gene signatures in single-cell datasets. UCell computes scores using the Mann-Whitney U statistic, which ensures robustness against variations in dataset size and heterogeneity. We created five gene signatures based on the top 50 differentially expressed genes between mast cell clusters in our dataset, and UCell was used to assign a score to each cell accordingly.

Cell signatures (**Fig. 3**) were generated by selecting the top 50 genes expressed in each cluster. The Seurat AddModuleScore function was then used to compute the average expression levels of these gene signatures (or modules) and assign a score to each cell, allowing for the evaluation of specific biological processes or pathways active within different cell populations. For each gene in the signatures, Seurat selects control genes with similar expression characteristics (matched by average expression and variance) to create a control gene set, used to reduce technical or dataset-specific noise. The function calculates the average expression of the signature genes, subtracts the average expression of the control genes, and produces a module score for each cell, reflecting how active or enriched the gene module is in comparison to the background.

**Figure 3.**
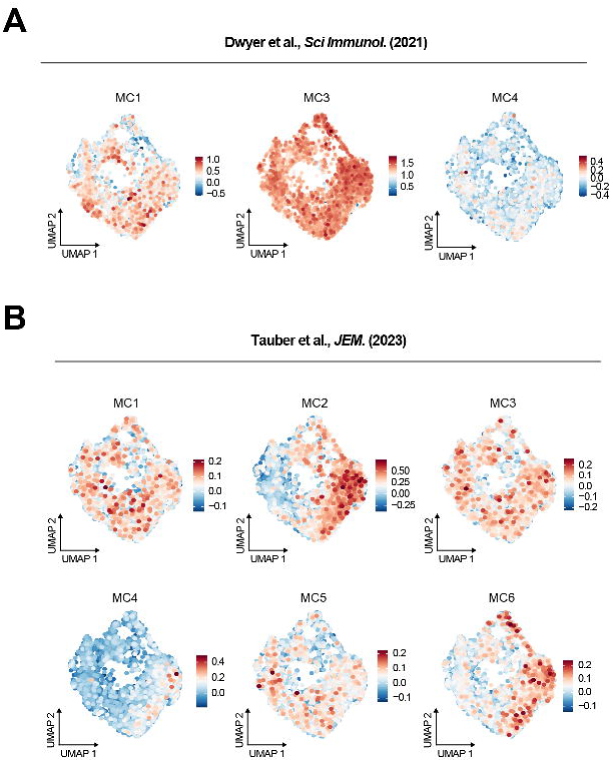
Mast cell subsets across single cell transcriptomic studies. Feature plots showing the expression of mast cell signatures described in Dwyer et al., *Sci Immunol* (2021) (A) and Tauber et al., *JEM* (2023) (B).

## Results

### Single cell RNA sequencing of healthy human colonic and rectal mast cells

In order to appropriately account for the heterogeneity of mast cells across the different microanatomical niches of the human gut, we decided to use dissected layers of the sigmoid colon, thereby obtaining individual mucosa, submucosa, and muscularis externa samples. Full thickness human gut samples can only be obtained during gut resection surgeries, which heavily limits the availability of such samples to study disease. Gut tissue biopsies, on the other hand, depend on less invasive and lengthy procedures for collection and are therefore more routinely available for the study of pathological conditions. We therefore included healthy rectal biopsies in our analysis to maximize the translational aspect of our data. We FACS-sorted live CD45^+^ cells from single cell suspensions obtained from healthy human gut tissues and generated our dataset using the 10x scRNA-seq protocol (**Fig. 1A**). Using the Uniform manifold approximation and projection (UMAP) dimensionality reduction technique, we clustered 137,087 cells at the CD45 level (**Fig. 1B**). We annotated immune subsets based on classical marker gene expression (**Fig. 1C**) and could identify plasma cells (*SDC1*), B cells (*CD19*), T cells (*CD3E*), myeloid cells (*CD14*), NK cells (*FCGR3A*), proliferating cells (*MKI67*), innate lymphoid cells (*KIT* and *RORC*), and mast cells (*KIT* and *CPA3*). Our dataset also contained contaminating non-immune cell subsets including fibroblasts (*COL1A1*), epithelial cells (*EPCAM*), endothelial cells (*CDH5*), and enteric glial cells (*SOX10*). The mast cell cluster comprised 4,148 cells. In comparison to other immune and non-immune cell types, mast cells expressed high levels of genes involved in mast cell development and function, such as the receptor for stem cell factor *KIT*, needed for mast cell maturation and survival, the beta subunit of the high affinity IgE receptor (*MS4A2*), involved in IgE-mediated mast cell activation, transcription factor GATA2, mast cell proteases tryptase (*TPSB2, TPSAB1*), carboxypeptidase A3 (*CPA3*), cathepsin G (*CTSG*), and chymase (*CMA1*), and hematopoietic prostaglandin D synthase (*HPGDS*), 15-hydroxyprostaglandin dehydrogenase (*HPGD*), and leukotriene C4 synthase (*LTC4S*), involved in eicosanoid metabolism. Mast cells also upregulated the receptor for IL-33 (*IL1RL1*), which potentiates mast cell responses^22^, histidine decarboxylase (*HDC*), involved in histamine biosynthesis, *SLC18A2*, a solute transporter involved in loading histamine and serotonin into secretory vesicles^23^, and amphiregulin (*AREG*), a secreted mediator member of the epidermal growth factor family involved in tissue regeneration^24^ (**Fig. 1D**). As previously described, *RHEX* expression was highly restricted to mast cells and has been described as a negative regulator of SCF/KIT signaling in skin mast cells^25^. Galectin 3 (*LGALS3*) is a negative regulator of IgE-mediated mast cell degranulation^26^. Tetraspanin CD9 is strongly expressed by glial cells in the gut, but also shows some level of expression in mast cells. CD9 has effects on mast cell chemotaxis, by acting as an alternate IL-16 receptor in human and mouse mast cells^27^ and on non-IgE mediated mast cell degranulation^28^. Gut mast cells also expressed high levels of Colony Stimulating Factor 1 (*CSF1*), indicating its potential role in supporting myeloid cells in the human gut^29^. Human gut mast cells also expressed high levels of the von-Willebrand factor A domain-containing protein 5A (*VWA5A*), also expressed by skin mast cells^30,31^, lamin A/C (*LMNA*), a protein of the nuclear lamina, and annexin A1 (*ANXA1*), previously shown in human lung mast cells^32^. ANXA1 is an anti-inflammatory molecule upregulated by glucocorticoids^33^. ANXA1 inhibits mast cell degranulation in mouse colonic mast cells and in human mast cells through the Formyl Peptide Receptor 2 (FPR2)^34,35^. Perilipin 2 (*PLIN2*) participates in the storage of neutral lipids in lipid droplets, which have been shown to develop in mast cells^36^ and to be involved in eicosanoid production. CAPG is an actin regulatory protein, and acts by capping the barbed ends of actin filaments, which is a critical step in the regulation of actin-based cell motility and function. CAPG is required for receptor-mediated ruffling, phagocytosis, and vesicle rocketing in macrophages^37^, suggesting that it could play a similar role in mast cells. *ADCYAP1* is a pleiotropic neuropeptide encoding the Pituitary Adenylate Cyclase-activating Polypeptide (PACAP). Expression of PACAP was previously shown in human mast cells in skin, colon, and small intestine^38^. SERPINB1 is a protease inhibitor which targets cathepsin G and chymase as well as neutrophil-derived proteases and protects mast cells from protease-induced cell death^39^. LEO1, a subunit of the conserved RNA polymerase-associated factor 1 complex, is required for the survival of yeast cells and human fibroblasts during quiescence^40,41^. The GTPase RAB27B regulates granule dynamics and secretion in human bone marrow-derived mast cells^42^. GPR65 is a pH-sensing receptor. While its role in controlling eosinophil survival to acidification was demonstrated, this was not the case for mast cells and the role of GPR65 in mast cells remains unclear^43^.

We then performed SCENIC analysis to identify enriched regulons, i.e., transcription factors and their target genes, among mast cells (**Fig. 1E**). The top 25 regulons in mast cells included genes required for mast cell progenitor development (*TAL1*^44^), mast cell development (*MITF*^45^, *GATA1*^46^, *GATA2*^47^), and mast cell function (*MITF*^48^, *GATA1*^49–51^, *GATA2*^48–53^*, EGR1*^54–56^*, NFE2L2*^57^*, BATF*^58^*, RARA*^59^). *BHLHE40* was recently identified as a top mast cell enhancer, but its function remains unknown in human mast cells^60^. Other identified genes have been shown to be expressed in human mast cells *ex vivo* (*NFE2L3*^30,61^, *MEIS2*^30,62^, *HOXB2*^30^) or *in vitro* (*EGR3*^30^, *RFX2*^30^, *SOX13*^63^), but their function in mast cells remains unknown. Our results identified four genes (*KLF6, MSX2, SOX4, ERG*) which have not previously been described in mast cells. Kruppel like factor 6 (*KLF6*) is a tumor suppressor gene belonging to the family of zinc finger transcription factors. KLF6 binds to the aryl hydrocarbon receptor (AhR), as part of the non-canonical signaling pathway^64^. AhR has been involved in the production of IL-6, IL-13, and IL-17 by mast cells^65^. KLF6 plays a pro-inflammatory role in intestinal macrophages, by upregulating NF-kB and suppressing STAT3 signaling in myeloid cells^66^. Msh homeobox 2 (*MSX2*) is a transcriptional repressor involved in craniofacial morphogenesis^67^ and mediate entry of human pluripotent stem cells into mesendoderm^68^. SOX4 belongs to the SRY-related HMG-box family of transcription factors. SOX proteins play important roles in cell fate specific specification and differentiation^69^. As the activity of SOX proteins depends on their binding partners, it will be important to determine the binding network of SOX4 in mast cells to delineate its functions. ETS-related gene (*ERG*) is an endothelial-specific transcription factor, as indicated by its high expression in endothelial cells in our dataset, involved in maintaining cell quiescence and homeostasis. ERG has been suggested to modulate microtubule dynamics^70^ as well as endothelial cell migration^71^ and to promote stiffness in cirrhotic liver endothelial cells in response to inflammatory cues^72^. ERG could therefore act in response to mechanical and immune signals and regulate the mast cell cytoskeleton, which plays an essential role in mast cell function^73^. In addition, human gut mast cells expressed regulons linked to the AP-1 (*JUN*, *FOSB*) and NF-κB (*NFKB1, NFKB2, RELB*) signaling pathways, which are more broadly expressed across other identified cell types.

### The human gut harbors five transcriptionally distinct mast cell subsets

To better understand mast cell heterogeneity in the human gut and identify potential mast cell subsets, we performed a subclustering of our initial mast cell population (**Fig. 2A**). The top 50 differentially expressed genes (DEGs) characterizing each mast cell subset, hereafter named MC1–5, is shown in **Fig. 2B**. The total list of DEGs of each mast cell subset, with relevant putative function and associated literature, is provided in **Table S2**. We first compared cytokine and chemokine expression in mast cell subsets. Distinct mast cell subsets could be distinguished based on patterns of cytokine and chemokine expression (**Fig. 2C**). MC1 showed high differential expression of *IL1B* and *IL13*, and of chemokines *CCL20* and *CXCL1*. CCL20 attracts lymphocytes and dendritic cells by binding to CCR6, while CXCL1 is involved in neutrophil chemotaxis. MC2 expressed high levels of *CSF1* and *LIF*, but also of *IL18*. LIF is a pleiotropic cytokine which can promote mast cell growth *in vitro* via its effects on fibroblasts^74,75^. IL-18 was suggested to act as an autocrine factor involved in mucosal mast cell development^76^. The MC3 subset expressed high levels of chemokines *CCL4* and *CCL4L2*, involved in T cell and myeloid cell recruitment, and *CXCL16*, which attracts T and NKT cells. Compared to other subsets, MC5 expressed high levels of *TGFB1*. TGF-β inhibits IgE- and IL-33-mediated mast cell activation in mouse and human mast cells, including TNF-α production^77,78^. Incidentally, MC5 also expressed the lowest level of *TNF* in our data. Whether TGF-β1 produced by mast cells is activated and is able to act in an autocrine fashion to control TNF-α production needs further investigation. MC5 and MC1 also expressed *TNFSF10*, encoding TRAIL, a pro-apoptotic cytokine. MC4 did not exhibit a marked cytokine signature, but showed expression of *CCL2*, a monocyte chemoattractant, also expressed by MC3, and *TNF*, also present in MC2.

To get further insight into the functions of mast cell subsets in the gut, we performed gene set enrichment analysis (**Fig. 2D**). Mast cell subsets upregulated certain shared pathways linked to mast cell function, such as the regulation of cysteine-type endopeptidase activity, response to TNF-α, or linked to apoptotic or transcriptional processes. MC1 were enriched in genes involved in the unfolded protein response (UPR), a state of endoplasmic reticulum stress linked to the accumulation of topologically incorrect proteins. This was in line with the high expression of heat shock proteins in the MC1 subset (**Fig. 2B**, **Table S2**). While molecular chaperones are often dismissed as a byproduct of stressful cell isolations procedures prior to library preparation and excluded from downstream analyses, the UPR exerts effects on immune cell survival and metabolism, but also on immune cell function and fate^79^. Of relevance to mast cell function, the UPR has been involved in prostaglandin biosynthesis^80^. The MC1 subset also showed enrichment in genes involved in cytokine response, antigen processing and presentation, and canonical NF-κB signal transduction. Indeed, we found that compared to other subsets, MC1 upregulated cytokine receptor genes, as well as genes involved in antigen presentation and co-stimulation (**Fig. 2E, F**). MC1 exhibited the highest levels of *IL1RL1*, *IL18R1*, and *IL4R* among gut mast cell subsets. MC1 also upregulated genes encoding for IL-2, IL-6, interferon type 1 and type 2, and TGF-β receptors. Mast cells have previously been shown to act as antigen presenting cells^81^. Here, we show that MC1 upregulate class II (*HLA-DRA*, *HLA-DRB1*, *HLA-DPA1*, *HLA-DPB1*) and class I (*HLA-A*, *HLA-B*, *HLA-C*) MHC molecules. MC1 also upregulated the non-classical MHC class I molecules, *HLA-E*, linked to NK cell activation. While MC1 expressed low levels of co-stimulatory molecules CD80, CD86, CD70, and CD40, they expressed higher levels of ICAM1, CD83, and CD274 (PDL-1). ICAM1 plays an important role as a co-stimulatory ligand during MHC-I-restricted antigen presentation^82^, while CD83 and CD274 (PDL-1) act as negative co-stimulatory molecules, suggesting a potential role of mast cells in negative co-stimulation in the human gut. MC1 also upregulated cytoskeleton-associated genes, both linked to microtubule (*TUBA1A*, *TUBA1B*, *TUBB4B*, *TUBB6*) or microfilament (*ACTG1*, *CDC42EP3*, *VASP*, *SAMSN1*, *EFHD2*) function. Mast cell activation and degranulation, which trigger changes in cell morphology, substrate adhesion, exocytosis, and migration, rely heavily on cytoskeletal processes^83^. Of note, lamin A/C was highly upregulated by MC1. Interestingly, lamin A participates in the formation and the stability of the immunological synapse between T cells and antigen presenting cells^84^. A similar role was found for transgellin 2 (*TAGLN2*), also upregulated in MC1^85^. The expression of synapse-stabilizing molecules could therefore support antigen presentation efficacy by MC1.

Differential expression analysis between mast cell clusters in the human gut also revealed that MC3 and MC4 were enriched in MC_TC_ signature genes (*HPGD, CMA1, CTSG*; **Fig. 2B**, **Fig. 2G**). Conversely, MC1 expressed low levels of MC_TC_ genes but upregulated histidine decarboxylase (*HDC*), suggesting that MC1 exhibit an MC_T_ phenotype (**Fig. 2B**). The MC_TC_ signature also included other genes, such as galectin 3 (*LGALS3*), serglycin (*SRGN*), neurotrimin (*NTM*), glutaredoxin (*GLRX*), metallothionein (*MT2A*), amphiregulin (*AREG*), and perilipin 2 (*PLIN2*). Expression of neurotrimin has been shown in skin mast cells at the protein level^31^ and has been suggested to act as a neural adhesion molecule and allow mast cells to localize close to nerves^86^. Interestingly, MC3 and MC4 differed by the expression of genes involved in prostaglandin metabolism, with MC3 expressing high levels of prostaglandin endoperoxide-synthase (*PTGS2*), more commonly called cyclooxygenase-2, while MC4 expressed the highest levels of 15-hydroxyprostaglandin dehydrogenase (*HPGD*) among mast cell subsets. PTGS2 catalyzes the first step of the prostaglandin synthesis pathway by converting arachidonic acid into prostaglandin H2, which can then be converted into prostaglandins (PGD2, PGE2, PGF2α), prostacyclin (PGI2), or thromboxane A2 by tissue-specific isomerases^87^. On the other hand, HPGD catalyzes the first step of the catabolic pathway of prostaglandins and therefore contributes to prostaglandin inactivation^88^. These observations highlight a differential role of MC3 and MC4 in prostaglandin signaling in the gut. MC3 was also characterized by its high expression of VEGFA, involved in angiogenesis, chemokines CCL4 and CCL4L2, which attract myeloid cells and T cells, amphiregulin, and the urokinase receptor (*PLAUR*), both involved in wound healing^24,89^.

We further observed that the MC1 and MC_TC_ transcriptional signatures were expressed along gradients, with the MC2 subset expressing low levels of each (**Fig. 2B**, **Fig. 2H**). This is in line with observations made by Dwyer et al. in nasal polyp mast cells. In our case, MC2 also exhibited a specific transcriptional signature, enriched in AP-1 (*JUN*, *FOSB*, *FOS*, *JUNB*) as well as other transcription factors (*EGR1*, *IER2*, *KLF2*, *KLF4*, *SOX4*, *IRF1*, *HES1*, *KLF6*, *EGR3*, *ATF3*) (**Fig. 2B**, **Fig. 2I**). MC2 was also enriched in genes involved in cell survival and quiescence (*TSC22D3*, *BTG1*, *BTG2*), the inhibition of cell signaling (*DUSP1*, *PPP1R15A*, *RGS1*, *PPP1R10*), and mRNA decay (*ZFP36*, *ZFP36L2*). Tristetraprolin (ZFP36), and other members of its family, ZFP36L1, and ZFP36L2, in humans, bind to adenine uridine-rich elements (AREs) in the 3’-UTRs of specific mRNAs and lead to target mRNA decay. This is notably the case of cytokines such as TNF-α. MC2, followed by MC4, expressed the highest levels of ZFP36 and ZFP36L2, while ZFP36L1 expression was low among all mast cell subsets (**Fig. 2J**). MC1 expressed ARE-containing genes *CCL20*, *CXCL1*, *CXCL8*, *HIF1A*, and *IL1B*, while MC3 expressed *PTGS2*, and *VEGFA*^90^ (**Fig. 2K**). Interestingly, we observed that MC2, which shows an intermediate state between MC1 and MC3 signatures, expressed lower levels of ARE-containing genes and exhibited an intermediate phenotype between MC1 and MC3. Whether this downregulation depends on tristetraprolin proteins requires further investigation.

As predicted by pathway analysis (**Fig. 2D**), the MC5 subset was enriched in genes involved in cholesterol metabolism and lipid particle clearance, namely apolipoproteins E (*APOE*) and C1 (*APOC1*) (**Fig. 2L**). MC5 expressed several genes associated with TGF-β signaling. *APOE*, *APOC1*, and *APOC2*, as well as *IL17RB*, were upregulated by TGF-β1 signaling in human M2 macrophages^91^. Olfactomedin 2 (*OLFM2*) and NGS1 were also shown to upregulate the TGF-β signaling pathway^92,93^. Additionally, MC5 expressed inhibitory receptors, GPR35 and CD52. GPR35 is a G-protein coupled receptor acting as a suppressor of mast cell degranulation, which mediates the effects of mast cell stabilizer disodium cromoglycate^94^. CD52 is a marker of antigen-activated T cells which bind the inhibitory receptor SIGLEC10^95^. It has been used as a molecular target in advanced systemic mastocytosis^96^. Of note, the MC5 gene signature was mainly identified in rectum cells, while colonic cells found in the MC5 cluster only expressed CD52 (**Fig. 2M**).

### Mast cell subsets across single cell transcriptomic studies

We next sought to compare our results with the mast cell subset gene signatures identified in previous human scRNA-seq studies focusing on mast cells. We thereby extracted the transcriptomic signatures identified in Dwyer et al.^15^ (**Fig. 3A**) and Tauber et al.^7^ (**Fig. 3B**) and projected them on our gut mast cell UMAP. Dwyer et al. described four mast cell populations in nasal polyps from chronic rhinosinusitis patients: MC1, expressing MC_TC_-associated genes (*CMA1*, *CTSG*, *PTGS2*), MC3 which did not express MC_TC_ genes, MC2, which showed an intermediate phenotype between MC1 and MC3, and MC4 which upregulated cell proliferation genes. Both MC1 and MC3 signatures were enriched in our dataset, however, this enrichment was not specific to a given mast cell subset (**Fig. 3A**). Using publicly available human transcriptomic datasets, Tauber et al. identified six mast cell clusters (MC1–6) distributed across 12 organs. The MC4 signature was not enriched in our dataset, while the MC1, MC3, and MC5 signatures showed a low and indiscriminate enrichment across gut mast cell subsets (**Fig. 3B**). The MC2 gene signature was enriched in gut MC1 cells. MC6 also showed low enrichment in MC_T_ gut subsets (MC1, MC2, MC5). Of note, MC6 was enriched in the lung, along with the MC1 subset, which were both associated with high *APOE* expression. Overall, we found a low correspondence between previously identified mast cell signatures and those identified in the human gut.

### Regulatory networks in human gut mast cell subsets

To gain insight into the regulatory networks directing the phenotype of mast cell subsets in the human gut, we performed SCENIC analysis on our previously characterized mast cell clusters (**Fig. 4A**). Distinct regulons were upregulated in each identified mast cell subset. MC1 upregulated regulons associated to the NF-κB pathway (*RELB, NFKB1, REL, NFKB2, RELA*), AP-1 transcription factors (*MAFG, JUND, FOSB*), and the UPR response (*XBP1, DDIT3, ATF4*). These regulons were upregulated to a lesser extent by MC2, which additionally exhibited a distinct signature composed of *SOX4, KLF2, EGR1, ATF3, ELK4, IRF1*, and AP-1 transcription factors *JUN* and *FOSB*. Interestingly, the two identified MC_TC_ subsets, MC3 and MC4, exhibited distinct regulatory signatures. MC3 showed a clear regulatory signature featuring *KLF7* and *KLF3*, *BATF*, *ZNF836*, *CREM*, *GLIS3*, *NR4A3*, and *CEBPD*. MC4 exhibited a smaller regulatory network composed of *CTCF*, *ELF2*, and *FOS*. MC5 showed a less obvious regulatory signature compared to other mast cell subsets but showed some enrichment of the *ELK3*, *ELF4*, *ZNF460*, and *SMAD1* regulons, which were associated to the MC1 regulatory signature and were enriched to a similar extent in the MC2 subset. Of note, the transcription factors identified for MC4 and MC5 via SCENIC analysis showed very little transcript expression. **Figure 4B** shows the expression of relevant transcription factors empirically selected based on regulon enrichment in gut mast cell subsets as well as high overall transcriptional expression. Based solely on expression level, MC1 and MC2 were enriched to similar extents in *ATF4*, *DDIT3*, and *GATA2*. *KLF2* was enriched in MC2, while MC3 showed high *BATF* expression. Interestingly, and in line with pathway analysis (**Fig. 2D**), MC1, MC3 and MC5 showed an upregulation of NF-κB genes *NFKB1* and *REL*, while MC2 and MC4 expressed high levels of *EGR1*, and the AP-1 transcription factors *JUN* and *FOS*. **Figure 4C** summarizes regulatory networks in human gut mast cell subsets.

**Figure 4.**
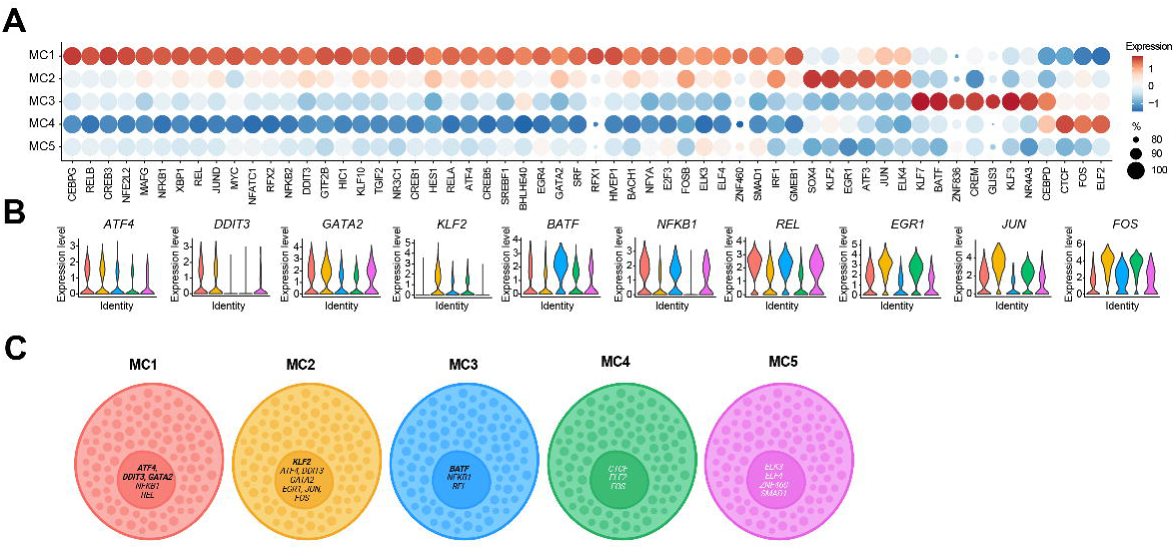
Regulatory networks in human gut mast cell subsets. (A) Heatmap showing the top regulons identified via single-cell regulatory network inference and clustering (SCENIC) analysis in human gut mast cell subsets. (B) Violin plots showing expression distribution of individual transcription factor genes associated to highly expressed regulons in mast cell subsets. (C) Graphical summary of the transcription factor network in mast cells subsets from the healthy rectum and colon based on transcription factor expression and SCENIC analysis.

### Mast cell subsets localize in specific layers of the healthy human gut

Finally, we investigated the anatomical distribution of distinct mast cell subsets within the human gut (**Fig. 5**). The colon mucosa was enriched in the MC1 and MC2 subsets and contained a small proportion of MC4 cells. The colon submucosa harbored the majority of MC4 cells, and a smaller proportion of MC2 and MC1 cells. MC3 was almost exclusively found in the colon muscularis, which also harbored a small proportion of other mast cell subsets. MC5 cells seemed enriched in colonic submucosa and muscularis as well as in rectal biopsies. However, mast cells exhibiting the full MC5 gene signature mainly localized in the rectum, and MC5 mast cells enriched in the colon submucosa and muscularis solely expressed CD52 (**Fig. 2M**).

**Figure 5.**
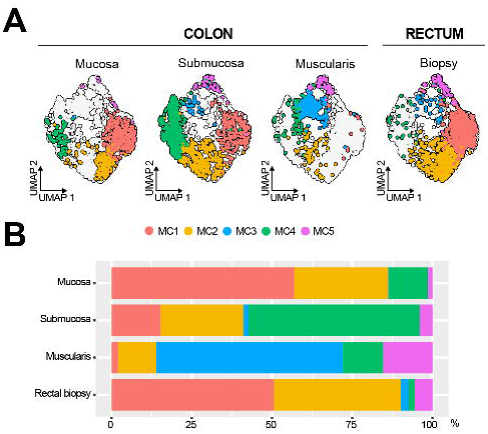
Mast cell subsets localize in specific layers of the healthy human gut. (A) UMAP of mast cells split by tissue layer (colon mucosa, submucosa, muscularis externa, rectal biopsies). (B) Barplot showing the distribution of mast cell subsets across colon layers and in rectal biopsies.

## Discussion

The field of mast cell biology is currently focused on unraveling the full spectrum of mast cell heterogeneity across tissues. Several attempts have clarified that mast cell complexity goes beyond the classical histological categorization of mucosal and connective tissue mast cells^7,15,60^. Here we provide the first comprehensive transcriptional analysis of human gut mast cells. We identified five distinct mast cells clusters (MC1-MC5) in the sigmoid colon and in rectal biopsies. MC1 was enriched in the mucosa of the colon and the rectum and was primed to interact with immune cells as it expressed high levels of cytokine and chemokine receptors, antigen presentation and co-stimulatory molecules, and cytoskeleton genes, suggestive of cell activation. MC3 and MC4 were mainly found in the muscularis and submucosa, respectively, and expressed a shared MC_TC_ transcriptional signature, including chymase (*CMA1*), 15-hydroxyprostaglandin dehydrogenase (*HPGD*), and cathepsin G (*CTSG*). MC3 and MC4 also upregulated the expression of neural adhesion marker neurotrimin (*NTM*), suggestive of a colocalization with neurons. MC3 and MC4 are likely to have different effects on prostaglandin availability in the colon, with MC3 upregulating *PTGS2*, linked to prostaglandin synthesis, and MC4 upregulating *HPGD*, linked to prostaglandin inactivation. Members of the prostaglandin family exert different effects on gastrointestinal function, including secretion and motility. Interestingly, fibroblasts from different layers of the colon expressed different prostaglandin isomerase genes (data not shown). This suggests that mast cells and fibroblasts collaborate to control gastrointestinal function via the regulation of prostaglandins in distinct layers of the human colon.

MC2 is an intermediate mast cell subtype found in the mucosa and submucosa, similar to the intermediate mast cell subset identified in nasal polyps^15^. While mucosal and connective tissue mast cells are thought to have different developmental origins, these results suggest that the local microenvironment also contributes in driving a specific transcriptional phenotype in mast cells. Contrastingly to the situation in the nose, however, this intermediate subset also exhibited a distinct transcriptional signature in the gut, composed of genes involved in quiescence, cell survival, and mRNA decay, and showed decreased cytokine and chemokine expression. We therefore hypothesize that the MC2 subset is a lowly active, quiescent mast cell subset that is capable of upregulating the gene signature of either mast cell effector subset (MC1, or MC3/MC4) in the right microenvironment. The existence of an intermediate mast cell phenotype also raises the question of the origin^97^ of these cells and whether connective tissue mast cells can differentiate from bone marrow-derived progenitors *in vivo*, as is the case in *in vitro* mast cell models. The fifth and last mast cell subset, MC5, was primarily located in the rectum and upregulated a gene signature suggestive of TGF-β signaling. Among these genes were several apoliproteins, lipid-binding proteins which participate in lipid metabolism, uptake, and transport. A specialized population of mast cells, termed intraepithelial mast cells, was found to populate the intestinal epithelium in mice. Intraepithelial mast cells were recently found to drive gasdermin C-mediated type 2 immunity in murine helminth infection^98^. In this context, intraepithelial mast cells exhibited a distinct transcriptional signature, characterized by *Mcpt1* expression, upregulated the apolipoprotein C4 (*Apoc4*) gene, and could be differentiated *in vitro* from bone marrow-derived progenitors after incubation with IL-9, SCF, and TGF-β1. Future work will have to determine whether human gut MC5 and murine intraepithelial mast cells are related subsets.

In this study, we also provide insight into the regulatory networks involved in the identified mast cell phenotypes in the human gut. The mucosal subset MC1 upregulated a large number of regulons, and could be associated with high expression of GATA2 and UPR-associated transcription factors ATF4 and DDIT3. This signature was expressed to a lesser extent by MC2, potentially indicating that MC2 derive from MC1 cells which become quiescent after upregulating KLF2, among others. MC3 and MC4 cells, which would have been classified as connective tissue mast cells based on their expression of chymase, showed distinct regulatory signatures, despite their phenotypic similarities. MC3 phenotype was associated with BATF, which has been associated with tissue residency and tissue repair phenotypes in human and mouse regulatory T cells^99–101^. Additionally, we showed an inverse relationship between NF-κ and AP-1 transcription factor expression among mast cell subsets, with mast cells in the mucosa and muscularis upregulating NF-κB regulons while submucosal subsets upregulated AP-1 transcription factors. Determining which environmental or functional signals are involved in the upregulation of specific regulatory networks in human mast cells will require further investigation.

Overall, our study expands our understanding of mast cell heterogeneity by characterizing five novel mast cell subsets associated with microanatomical niches in the human gut. Our study also provides further evidence of the existence of a transcriptional gradient in mast cell effector subsets. Importantly, while transcriptomic studies are an important tool which has successfully been used to progressively unravel mast cell heterogeneity, transcriptional observations relating to mast cell phenotypes will have to be validated at the protein level in mouse and in human. Mast cells play important roles in gastrointestinal diseases^102–104^, notably through their capacity to interact with the enteric nervous system^105^. How the five mast cell subsets identified in this study interact with the intestinal immune and nervous systems and how they contribute to gastrointestinal disease development will require further investigation.

## Supporting information

Supplemental table 1 - Reagents and media

Supplemental table 2 - DEG MC1 to MC5

## Author contributions

H.H. designed the study, conducted experiments, and wrote the manuscript. E.M. performed bioinformatic analysis of scRNA-seq data. N.S. and M.D. collected sigmoid colon samples. A.S., K.V., and T.V. provided intellectual input. G.B. led the project and revised the manuscript.

## Acknowledgements

This work is supported by an FWO grant (G077222N).

## References

1. Chen, E. et al. Inflamed Ulcerative Colitis Regions Associated With MRGPRX2-Mediated Mast Cell Degranulation and Cell Activation Modules, Defining a New Therapeutic Target. Gastroenterology 160, 1709–1724 (2021).

2. Van Remoortel, S. et al. Mrgprb2-dependent Mast Cell Activation Plays a Crucial Role in Acute Colitis. Cell. Mol. Gastroenterol. Hepatol. 18, 101391 (2024).

3. Galli, S. J., Gaudenzio, N. & Tsai, M. Mast Cells in Inflammation and Disease: Recent Progress and Ongoing Concerns. Annu. Rev. Immunol. 38, 49–77 (2020).

4. Wong, G. W. et al. Ancient origin of mast cells. Biochem. Biophys. Res. Commun. 451, 314–318 (2014).

5. Cavalcante, M. C. M. et al. Occurrence of Heparin in the Invertebrate *Styela plicata* (Tunicata) Is Restricted to Cell Layers Facing the Outside Environment. J. Biol. Chem. 275, 36189–36196 (2000).

6. Moon, T. C., Befus, A. D. & Kulka, M. Mast Cell Mediators: Their Differential Release and the Secretory Pathways Involved. Front. Immunol. 5, 569 (2014).

7. Tauber, M. et al. Landscape of mast cell populations across organs in mice and humans. J. Exp. Med. 220, e20230570 (2023).

8. Gentek, R. et al. Hemogenic Endothelial Fate Mapping Reveals Dual Developmental Origin of Mast Cells. Immunity 48, 1160–1171.e5 (2018).

9. Shimokawa, C. et al. Mast Cells Are Crucial for Induction of Group 2 Innate Lymphoid Cells and Clearance of Helminth Infections. Immunity 46, 863–874.e4 (2017).

10. Groschwitz, K. R. et al. Mast cells regulate homeostatic intestinal epithelial migration and barrier function by a chymase/Mcpt4-dependent mechanism. Proc. Natl. Acad. Sci. U. S. A. 106, 22381–22386 (2009).

11. Albert-Bayo, M. et al. Intestinal Mucosal Mast Cells: Key Modulators of Barrier Function and Homeostasis. Cells 8, (2019).

12. De Giovanni, M., et al. Mast cells help organize the Peyer’s patch niche for induction of IgA resonses. Sci. Immunol. 9, eadj7363 (2024).

13. Wood, J. D. Enteric neuroimmunophysiology and pathophysiology1,2. Gastroenterology 127, 635–657 (2004).

14. Sharkey, K. A. & Mawe, G. M. The enteric nervous system. Physiol. Rev. 103, 1487–1564 (2023).

15. Dwyer, D. F., et al. Human airway mast cells proliferate and acquire distinct inflammation-driven phenotypes during type 2 inflammation. Sci. Immunol. 6, eabb7221 (2021).

16. Young, M. D. & Behjati, S. SoupX removes ambient RNA contamination from droplet-based single-cell RNA sequencing data. GigaScience 9, giaa151 (2020).

17. Korsunsky, I. et al. Fast, sensitive and accurate integration of single-cell data with Harmony. Nat. Methods 16, 1289–1296 (2019).

18. McGinnis, C. S., Murrow, L. M. & Gartner, Z. J. DoubletFinder: Doublet Detection in Single-Cell RNA Sequencing Data Using Artificial Nearest Neighbors. Cell Syst. 8, 329–337.e4 (2019).

19. Aibar, S. et al. SCENIC: single-cell regulatory network inference and clustering. Nat. Methods 14, 1083–1086 (2017).

20. Kolberg, L., Raudvere, U., Kuzmin, I., Vilo, J. & Peterson, H. gprofiler2 -- an R package for gene list functional enrichment analysis and namespace conversion toolset g:Profiler. F1000Research 9, ELIXIR-709 (2020).

21. Andreatta, M. & Carmona, S. J. UCell: Robust and scalable single-cell gene signature scoring. Comput. Struct. Biotechnol. J. 19, 3796–3798 (2021).

22. Joulia, R., L’Faqihi, F.-E., Valitutti, S. & Espinosa, E. IL-33 fine tunes mast cell degranulation and chemokine production at the single-cell level. J. Allergy Clin. Immunol. 140, 497–509.e10 (2017).

23. Dwyer, D. F., Barrett, N. A. & Austen, K. F. Expression profiling of constitutive mast cells reveals a unique identity within the immune system. Nat. Immunol. 17, 878–887 (2016).

24. Zaiss, D. M. W., Gause, W. C., Osborne, L. C. & Artis, D. Emerging functions of amphiregulin in orchestrating immunity, inflammation, and tissue repair. Immunity 42, 216–226 (2015).

25. Franke, K., Bal, G., Li, Z., Zuberbier, T. & Babina, M. Clorfl86/RHEX Is a Negative Regulator of SCF/KIT Signaling in Human Skin Mast Cells. Cells 12, 1306 (2023).

26. Bambouskova, M. et al. New Regulatory Roles of Galectin-3 in High-Affinity IgE Receptor Signaling. Mol. Cell. Biol. 36, 1366–1382 (2016).

27. Qi, J. C. et al. Human and mouse mast cells use the tetraspanin CD9 as an alternate interleukin-16 receptor. Blood 107, 135–142 (2006).

28. Hálová, I. et al. Cross-talk between Tetraspanin CD9 and Transmembrane Adaptor Protein Non-T Cell Activation Linker (NTAL) in Mast Cell Activation and Chemotaxis*. J. Biol. Chem. 288, 9801–9814 (2013).

29. Yadav, S., Priya, A., Borade, D. R. & Agrawal-Rajput, R. Macrophage subsets and their role: co-relation with colony-stimulating factor-1 receptor and clinical relevance. Immunol. Res. 71, 130–152 (2023).

30. Motakis, E. et al. Redefinition of the human mast cell transcriptome by deep-CAGE sequencing. Blood 123, e58–e67 (2014).

31. Plum, T. et al. Human Mast Cell Proteome Reveals Unique Lineage, Putative Functions, and Structural Basis for Cell Ablation. Immunity 52, 404–416.e5 (2020).

32. Rönnberg, E. et al. Analysis of human lung mast cells by single cell RNA sequencing. Front. Immunol. 14, (2023).

33. Purvis, G. S. D., Solito, E. & Thiemermann, C. Annexin-A1: Therapeutic Potential in Microvascular Disease. Front. Immunol. 10, (2019).

34. Sinniah, A. et al. Endogenous Annexin-A1 Negatively Regulates Mast Cell-Mediated Allergic Reactions. Front. Pharmacol. 10, 1313 (2019).

35. Oliveira, M. P., Prates, J., Gimenes, A. D., Correa, S. G. & Oliani, S. M. Annexin A1 Mimetic Peptide Ac2-26 Modulates the Function of Murine Colonic and Human Mast Cells. Front. Immunol. 12, 689484 (2021).

36. Dichlberger, A. et al. Lipid body formation during maturation of human mast cells. J. Lipid Res. 52, 2198–2208 (2011).

37. Witke, W., Li, W., Kwiatkowski, D. J. & Southwick, F. S. Comparisons of CapG and gelsolin-null macrophages. J. Cell Biol. 154, 775–784 (2001).

38. Okragly, A. J. et al. Human mast cells release the migraine-inducing factor pituitary adenylate cyclase-activating polypeptide (PACAP). Cephalalgia Int. J. Headache 38, 1564–1574 (2018).

39. Burgener, S. S. et al. Granule Leakage Induces Cell-Intrinsic, Granzyme B-Mediated Apoptosis in Mast Cells. Front. Cell Dev. Biol. 9, 630166 (2021).

40. Oya, E. et al. Leo1 is essential for the dynamic regulation of heterochromatin and gene expression during cellular quiescence. Epigenetics Chromatin 12, 45 (2019).

41. Laurent, M., Cordeddu, L., Zahedi, Y. & Ekwall, K. LEO1 Is Required for Efficient Entry into Quiescence, Control of H3K9 Methylation and Gene Expression in Human Fibroblasts. Biomolecules 13, 1662 (2023).

42. Mizuno, K. et al. Rab27b regulates mast cell granule dynamics and secretion. Traffic Cph. Den. 8, 883–892 (2007).

43. Zhu, X., Mose, E., Hogan, S. P. & Zimmermann, N. Differential eosinophil and mast cell regulation: Mast cell viability and accumulation in inflammatory tissue are independent of proton-sensing receptor GPR65. Am. J. Physiol. - Gastrointest. Liver Physiol. 306, G974–G982 (2014).

44. Salmon, J. M. et al. Aberrant mast-cell differentiation in mice lacking the stem-cell leukemia gene. Blood 110, 3573–3581 (2007).

45. Takemoto, C. M., Yoon, Y.-J. & Fisher, D. E. The identification and functional characterization of a novel mast cell isoform of the microphthalmia-associated transcription factor. J. Biol. Chem. 277, 30244–30252 (2002).

46. Migliaccio, A. R. et al. GATA-1 as a regulator of mast cell differentiation revealed by the phenotype of the GATA-1low mouse mutant. J. Exp. Med. 197, 281– 296 (2003).

47. Ohmori, S. et al. GATA2 is critical for the maintenance of cellular identity in differentiated mast cells derived from mouse bone marrow. Blood 125, 3306–3315 (2015).

48. Li, Y. et al. The transcription factors GATA2 and microphthalmia-associated transcription factor regulate Hdc gene expression in mast cells and are required for IgE/mast cell-mediated anaphylaxis. J. Allergy Clin. Immunol. 142, 1173–1184 (2018).

49. McLeod, J. J. A., Baker, B. & Ryan, J. J. Mast cell production and response to IL-4 and IL-13. Cytokine 75, 57–61 (2015).

50. Inage, E. et al. Critical Roles for PU.1, GATA1, and GATA2 in the expression of human FcεRI on mast cells: PU.1 and GATA1 transactivate FCER1A, and GATA2 transactivates FCER1A and MS4A2. J. Immunol. Baltim. Md 1950 192, 3936–3946 (2014).

51. Ohneda, K., Ohmori, S. & Yamamoto, M. Mouse Tryptase Gene Expression is Coordinately Regulated by GATA1 and GATA2 in Bone Marrow-Derived Mast Cells. Int. J. Mol. Sci. 20, 4603 (2019).

52. Maeda, K., Nishiyama, C., Ogawa, H. & Okumura, K. GATA2 and Sp1 positively regulate the c-kit promoter in mast cells. J. Immunol. Baltim. Md 1950 185, 4252–4260 (2010).

53. Baba, Y. et al. GATA2 is a critical transactivator for the human IL1RL1/ST2 promoter in mast cells/basophils: opposing roles for GATA2 and GATA1 in human IL1RL1/ST2 gene expression. J. Biol. Chem. 287, 32689–32696 (2012).

54. Wang, H.-N. et al. Inhibition of c-Fos expression attenuates IgE-mediated mast cell activation and allergic inflammation by counteracting an inhibitory AP1/Egr1/IL-4 axis. J. Transl. Med. 19, 261 (2021).

55. Li, B. et al. The early growth response factor-1 contributes to interleukin-13 production by mast cells in response to stem cell factor stimulation. J. Immunotoxicol. 5, 163–171 (2008).

56. Li, B., Power, M. R. & Lin, T.-J. De novo synthesis of early growth response factor-1 is required for the full responsiveness of mast cells to produce TNF and IL-13 by IgE and antigen stimulation. Blood 107, 2814–2820 (2006).

57. Jadkauskaite, L. et al. Nuclear factor (erythroid-derived 2)-like-2 pathway modulates substance P-induced human mast cell activation and degranulation in the hair follicle. J. Allergy Clin. Immunol. 142, 1331–1333.e8 (2018).

58. Tomar, S. et al. IL-4-BATF signaling directly modulates IL-9 producing mucosal mast cell (MMC9) function in experimental food allergy. J. Allergy Clin. Immunol. 147, 280–295 (2021).

59. Babina, M. et al. Retinoic acid potentiates inflammatory cytokines in human mast cells: identification of mast cells as prominent constituents of the skin retinoid network. Mol. Cell. Endocrinol. 406, 49–59 (2015).

60. Cildir, G. et al. Genome-wide Analyses of Chromatin State in Human Mast Cells Reveal Molecular Drivers and Mediators of Allergic and Inflammatory Diseases. Immunity 51, 949–965.e6 (2019).

61. Saliba, J. et al. Loss of NFE2L3 protects against inflammation-induced colorectal cancer through modulation of the tumor microenvironment. Oncogene 41, 1563–1575 (2022).

62. Hamey, F. K. et al. Single-cell molecular profiling provides a high-resolution map of basophil and mast cell development. Allergy 76, 1731–1742 (2021).

63. Jayapal, M. et al. Genome-wide gene expression profiling of human mast cells stimulated by IgE or FcεRI-aggregation reveals a complex network of genes involved in inflammatory responses. BMC Genomics 7, 210 (2006).

64. Wilson, S. R., Joshi, A. D. & Elferink, C. J. The tumor suppressor Kruppel-like factor 6 is a novel aryl hydrocarbon receptor DNA binding partner. J. Pharmacol. Exp. Ther. 345, 419–429 (2013).

65. Sibilano, R., Pucillo, C. E. & Gri, G. Allergic responses and aryl hydrocarbon receptor novel pathway of mast cell activation. Mol. Immunol. 63, 69–73 (2015).

66. Goodman, W. A. et al. KLF6 contributes to myeloid cell plasticity in the pathogenesis of intestinal inflammation. Mucosal Immunol. 9, 1250–1262 (2016).

67. Takahashi, K. et al. Msx2 is a repressor of chondrogenic differentiation in migratory cranial neural crest cells. Dev. Dyn. Off. Publ. Am. Assoc. Anat. 222, 252– 262 (2001).

68. Wu, Q. et al. MSX2 mediates entry of human pluripotent stem cells into mesendoderm by simultaneously suppressing SOX2 and activating NODAL signaling. Cell Res. 25, 1314–1332 (2015).

69. Kamachi, Y. & Kondoh, H. Sox proteins: regulators of cell fate specification and differentiation. Development 140, 4129–4144 (2013).

70. Yuan, L. et al. RhoJ is an endothelial cell-restricted Rho GTPase that mediates vascular morphogenesis and is regulated by the transcription factor ERG. Blood 118, 1145–1153 (2011).

71. Birdsey, G. M. et al. The transcription factor Erg regulates expression of histone deacetylase 6 and multiple pathways involved in endothelial cell migration and angiogenesis. Blood 119, 894–903 (2012).

72. Selicean, S.-E. et al. Stiffness-induced modulation of ERG transcription factor in chronic liver disease. Npj Gut Liver 1, 7 (2024).

73. Lazki-Hagenbach, P., Klein, O. & Sagi-Eisenberg, R. The actin cytoskeleton and mast cell function. Curr. Opin. Immunol. 72, 27–33 (2021).

74. Hiragun, T. et al. Leukemia inhibitory factor enhances mast cell growth in a mast cell/fibroblast co-culture system through stat3 signaling pathway of fibroblasts. FEBS Lett. 487, 219–223 (2000).

75. Gyotoku, E. et al. The IL-6 family cytokines, interleukin-6, interleukin-11, oncostatin M, and leukemia inhibitory factor, enhance mast cell growth through fibroblast-dependent pathway in mice. Arch. Dermatol. Res. 293, 508–514 (2001).

76. Sandersa, N. L. et al. Interleukin-18 has an Important Role in Differentiation and Maturation of Mucosal Mast Cells. J. Mucosal Immunol. Res. 2, 109 (2018).

77. Ndaw, V. S. et al. TGF-β1 Suppresses IL-33–Induced Mast Cell Function. J. Immunol. 199, 866–873 (2017).

78. Fernando, J. et al. Genotype-dependent Effects of TGFβ1 on Mast Cell Function: Targeting the Stat5 Pathway. J. Immunol. Baltim. Md 1950 191, 10.4049/jimmunol.1202723 (2013).

79. Di Conza, G., Ho, P.-C., Cubillos-Ruiz, J. R. & Huang, S. C.-C. Control of immune cell function by the unfolded protein response. Nat. Rev. Immunol. 23, 546– 562 (2023).

80. Chopra, S. et al. IRE1α-XBP1 signaling in leukocytes controls prostaglandin biosynthesis and pain. Science 365, eaau6499 (2019).

81. Galli, S. J. & Gaudenzio, N. Human mast cells as antigen-presenting cells: When is this role important in vivo? J. Allergy Clin. Immunol. 141, 92–93 (2018).

82. Lebedeva, T., Dustin, M. L. & Sykulev, Y. ICAM-1 co-stimulates target cells to facilitate antigen presentation. Curr. Opin. Immunol. 17, 251–258 (2005).

83. Dráber, P., Sulimenko, V. & Dráberová, E. Cytoskeleton in Mast Cell Signaling. Front. Immunol. 3, (2012).

84. González-Granado, J. M. et al. Nuclear envelope lamin-A couples actin dynamics with immunological synapse architecture and T cell activation. Sci. Signal. 7, ra37 (2014).

85. Na, B.-R. et al. TAGLN2 regulates T cell activation by stabilizing the actin cytoskeleton at the immunological synapse. J. Cell Biol. 209, 143–162 (2015).

86. Babina, M., Franke, K. & Bal, G. How “Neuronal” Are Human Skin Mast Cells? Int. J. Mol. Sci. 23, 10871 (2022).

87. Ricciotti, E. & FitzGerald, G. A. Prostaglandins and Inflammation. Arterioscler. Thromb. Vasc. Biol. 31, 986–1000 (2011).

88. Sun, C.-C. et al. Recent advances in studies of 15-PGDH as a key enzyme for the degradation of prostaglandins. Int. Immunopharmacol. 101, 108176 (2021).

89. Gonias, S. L. Plasminogen activator receptor assemblies in cell signaling, innate immunity, and inflammation. Am. J. Physiol. - Cell Physiol. 321, C721–C734 (2021).

90. Mukherjee, N. et al. Global target mRNA specification and regulation by the RNA-binding protein ZFP36. Genome Biol. 15, R12 (2014).

91. Gratchev, A. et al. Activation of a TGF-β-Specific Multistep Gene Expression Program in Mature Macrophages Requires Glucocorticoid-Mediated Surface Expression of TGF-β Receptor II1. J. Immunol. 180, 6553–6565 (2008).

92. Tu, M. et al. NSG1 promotes glycolytic metabolism to enhance Esophageal squamous cell carcinoma EMT process by upregulating TGF-β. Cell Death Discov. 9, 1–11 (2023).

93. Tang, Y. et al. OLFM2 promotes epithelial-mesenchymal transition, migration, and invasion in colorectal cancer through the TGF-β/Smad signaling pathway. BMC Cancer 24, 204 (2024).

94. Oka, M. et al. Suppression of Mast Cell Activation by GPR35: GPR35 Is a Primary Target of Disodium Cromoglycate. J. Pharmacol. Exp. Ther. 389, 76–86 (2024).

95. Bandala-Sanchez, E. et al. T cell regulation mediated by interaction of soluble CD52 with the inhibitory receptor Siglec-10. Nat. Immunol. 14, 741–748 (2013).

96. Hoermann, G. et al. CD52 is a molecular target in advanced systemic mastocytosis. FASEB J. Off. Publ. Fed. Am. Soc. Exp. Biol. 28, 3540–3551 (2014).

97. St. John, A. L., Rathore, A. P. S. & Ginhoux, F. New perspectives on the origins and heterogeneity of mast cells. Nat. Rev. Immunol. 23, 55–68 (2023).

98. Yang, L. et al. Intraepithelial mast cells drive gasdermin C-mediated type 2 immunity. Immunity 0, (2024).

99. Delacher, M. et al. Single-cell chromatin accessibility landscape identifies tissue repair program in human regulatory T cells. Immunity 54, 702–720.e17 (2021).

100. Delacher, M. et al. Precursors for Nonlymphoid-Tissue Treg Cells Reside in Secondary Lymphoid Organs and Are Programmed by the Transcription Factor BATF. Immunity 52, 295–312.e11 (2020).

101. Burton, O. T. et al. The tissue-resident regulatory T cell pool is shaped by transient multi-tissue migration and a conserved residency program. Immunity 57, 1586–1602.e10 (2024).

102. Molfetta, R. & Paolini, R. The Controversial Role of Intestinal Mast Cells in Colon Cancer. Cells 12, 459 (2023).

103. Hamilton, M. J. Gastrointestinal Disease in Mastocytosis. Immunol. Allergy Clin. North Am. 43, 711–722 (2023).

104. Hamilton, M. J., Frei, S. M. & Stevens, R. L. The Multifaceted Mast Cell in Inflammatory Bowel Disease. Inflamm. Bowel Dis. 20, 2364–2378 (2014).

105. Hussein, H., Van Remoortel, S. & Boeckxstaens, G. E. Irritable bowel syndrome: When food is a pain in the gut. Immunol. Rev. 326, 102–116 (2024).

